# Reliability of transcranial magnetic stimulation-evoked responses on knee extensor muscles during cycling

**DOI:** 10.1101/2024.02.12.579935

**Authors:** Jenny Zhang, Zachary J. McClean, Neda Khaledi, Sophie-Jayne Morgan, Guillaume Y. Millet, Saied Jalal Aboodarda

## Abstract

Transcranial magnetic stimulation (TMS) measures the excitability and inhibition of corticomotor networks. Despite its task-specificity, few studies have used TMS during dynamic movements and the reliability of TMS-derived measures has not been assessed during cycling. This study aimed to evaluate the reliability of motor evoked potentials (MEP) and short- and long-interval intracortical inhibition (SICI and LICI) on vastus lateralis and rectus femoris muscle activity during a fatiguing single-leg cycling task. Nine healthy adults (2 females) performed two identical sessions of counterweighted single-leg cycling at 60% peak power output until failure. Five single-pulses and five short- and long-interval paired pulses delivered to the motor cortex, and two maximal femoral nerve stimulations [maximal M-wave (M_max_)], were delivered during two baseline cycling bouts (unfatigued) and every 5 min throughout cycling (fatigued). When comparing both baseline bouts within the same session, MEP·M_max_^-1^ and LICI (both ICC: >0.9) were rated excellent while SICI was rated good (ICC: 0.7-0.9). At baseline between sessions, in the vastus lateralis, M_max_ (ICC: >0.9) and MEP·M_max_^-1^ (ICC: 0.7) demonstrated good reliability, LICI was moderate (ICC: 0.5), and SICI was poor (ICC: 0.3). Across the fatiguing task, M_max_ demonstrated excellent reliability (ICC >0.8), MEP·M_max_^-1^ ranged good to excellent (ICC: 0.7-0.9), LICI was moderate to excellent (ICC: 0.5-0.9), and SICI remained poorly reliable (ICC: 0.3-0.6). Overall, these results corroborate the cruciality of retaining mode-specific testing measurements and suggest that during cycling, M_max_, MEP·M_max_^-1^, and LICI measures are reliable whereas SICI, although less reliable across days, can be reliable within the same session.

## INTRODUCTION

Transcranial magnetic stimulation (TMS) is a non-invasive, widely used technique whereby descending neural volleys are induced atop the primary motor cortex to produce short-latency electromyography (EMG) signals, termed motor evoked potential (MEP), in the muscles of interest. The area and/or amplitude of MEP and the ensuing EMG silence [termed silent period (SP)] are measures of the excitation and inhibition of the corticospinal tract, respectively (Gandevia *et al*., 1996; Taylor *et al*., 2006). Utilization paired-pulse TMS paradigms can also help characterize the efficiency of GABA-mediated cortical inhibitory circuitries. By altering the interstimulus interval and observing the effect of a conditioning stimulus on the size of a subsequent test stimulus (i.e., MEP), short- and long-interval intracortical inhibition (SICI and LICI, respectively) can be quantified (Valls-Solé *et al*., 1992; Kujirai *et al*., 1993). These metrics can be assessed before, during, and after an exercise task as a method to explore the responsiveness of the corticomotor pathway (Taylor *et al*., 1996; Di Lazzaro *et al*., 1999; Taylor & Gandevia, 2004; Martin *et al*., 2008; McNeil *et al*., 2009; Hunter *et al*., 2016).

One phenomenon that can modulate corticospinal responsiveness is exercise-induced fatigue (McNeil *et al*., 2009; Hunter *et al*., 2016; Temesi *et al*., 2019) and its associated alterations in sensory feedback and motor feedforward discharges (Sidhu *et al*., 2017; Weavil & Amann, 2018; Azevedo *et al*., 2022). Numerous studies have used TMS to explore corticospinal responses to fatiguing exercise; however, such neurophysiological assessments are often measured during single-joint isometric contractions, whereas the effects of the exercise under investigation may have been whole body dynamic exercises such as cycling (Thomas *et al*., 2015, 2016), running (Temesi *et al*., 2014), or any other types of activities (Brownstein *et al*., 2017; Thomas *et al*., 2017). The ability to measure neurophysiological responses during, as opposed to only pre- and post-, a locomotor task is crucial for two primary reasons: firstly, this allows the measurements to be taken across an exercise task without interruptions required for isometric contractions, so that response kinetics can be evaluated more accurately and secondly, this allows a better understanding of corticomotoneuronal modulations during whole-body and/or large muscle mass dynamic movements which are biomechanically and physiologically distinct from single-limb isometric contractions. Various aspects of these distinctions can include muscle ischemia in isometric contractions which can modulate activity of group III/IV afferent fibres (Sadamoto *et al*., 1983; Sjogaard *et al*., 1988), greater cardiorespiratory demand from a larger volume of active muscles during whole-body movements (Andersen & Saltin, 1985), and the contribution of muscle spindles and/or golgi tendon organs across different contraction modes (Nelson & Hutton, 1985). To avoid the confounding effect of these examples, it is crucial that the exercise mode be retained between the task and the measurements.

To our knowledge, few studies have measured the responsiveness of corticospinal circuitries *during* leg cycling. These experiments demonstrated that unlike isometric single-joint exercises where MEP generally increases (Gandevia *et al*., 1996; Temesi *et al*., 2019), corticospinal excitability tends not to change throughout fatiguing exercise (Sidhu *et al*., 2012*a*; Weavil *et al*., 2016; Sidhu *et al*., 2017), while the excitability of intracortical inhibitory networks has been shown to both increase (Sidhu *et al*., 2013) and decrease during cycling (Sidhu *et al*., 2018). The equivocal observations of TMS-induced responses during isometric and dynamic contractions highlights the task-dependent nature of these responses that can vary based on numerous factors, including muscle contraction patterns. Despite the importance of this observation, the usage of TMS during leg cycling is a relatively novel and uncommon technique. Furthermore, although the reliability of TMS-evoked measures [i.e., MEP] has been reported during fatiguing isometric contractions (Todd *et al*., 2007; Doyle-Baker *et al*., 2018) and once during single-joint dynamic contractions (Tallent *et al*., 2012), as recently pointed out by Kesar et al. (2018), there is a lack of adequate test-retest reliability reporting of TMS measures in the lower limb muscles during fatiguing cycling contractions. Additionally, no prior study has reported the reliability of paired pulse paradigms to measure intracortical inhibition (i.e., SICI and LICI) during leg cycling, which may be useful to further delineate the excitation and inhibition along the corticospinal pathway. Characterizing day-to-day reliability of these measures can help delineate meaningful changes from inherent variability.

Therefore, the present study aimed to assess the reliability of single- and paired-pulse TMS paradigms evoked onto the knee extensor muscles during a fatiguing single-leg cycling task. This experiment was conducted as a part of a more comprehensive study measuring the effect of pain on contralateral limb corticomotor responses; thereby, single-leg cycling was used as the exercise framework herein.

## METHODS

### Participants

Nine healthy, recreationally active individuals volunteered for this study (females = 2, age = 30 ± 7 years, height = 176.5 ± 6.6 cm, weight = 78.9 ± 23.7 kg). All participants were non-smokers, free from any neurological, metabolic, cardiovascular, and/or mood disorders, and did not have a history of a lower limb injury within the past 6 months that could impair their ability to cycle or produce a maximal isometric contraction. All participants cleared the Physical Activity Readiness Questionnaire Plus and TMS safety checklist (Rossi *et al*., 2011), and provided their written informed consent prior to starting participation in the study. The study was approved by the Conjoint Health Research Ethics Board of the University of Calgary (REB21-0264).

### Experimental protocol

Participants attended the laboratory for three visits. All sessions consisted of single-leg cycling using the dominant (right) leg on a custom-built semi-recumbent ergometer (Doyle-Baker *et al*., 2018) with a custom 10-kg counterweight fitted in the contralateral pedal (Aboodarda *et al*., 2020).

### Ramp-incremental test

In the first visit, participants completed a single-leg maximal cycling test consisting of a 4 min baseline at 20 W followed immediately by a smooth 10 W·min^-1^ ramp (1 W/6 s) until they were unable to maintain a cadence above 60 rpm for more than 5 s despite strong verbal encouragement (i.e., task failure). Following the ramp and after a 15 min rest period, participants performed a 12 min heavy-intensity step transition at a power output corresponding to 60% of peak power output (PO_peak_) to ensure stabilization of oxygen uptake and to familiarize participants with the cycling intensity for future sessions. This intensity was prescribed based on two prior studies in our lab using the same counterweighted single-leg cycling setup, one of which is currently under review, wherein the power output corresponding to the heavy-intensity domain (i.e., between the gas exchange threshold and the maximal metabolic steady state) was approximately 60% of PO_peak_ from a single-leg ramp test of the same ramp rate (Zhang *et al*., 2021).

### Experimental sessions

The next two visits were identical, performed during the same time of day, and spaced apart by at least 72 h. Upon arrival, participants were first prepped for corticospinal measurements, after which muscle compound action potential (M_max_), TMS hotspot, active motor threshold (aMT), and SICI and LICI intensities were determined. Then, participants performed two 2 min baseline cycling bouts separated by at least 5 min rest to allow complete recovery during which MEP, M_max_, SICI and LICI were elicited. Following baseline bouts, participants performed the experimental trial consisting of constant-PO single-leg cycling at 60% PO_peak_ to task failure. Every 5 min, the cycle ergometer pedals were locked and within 1 s, participants performed a maximal and submaximal isometric contraction with stimuli before immediately resuming cycling (approximately 15 s); however, these results are not reported in the present study in the interest of reliability. During cycling, participants were instructed to maintain a consistent cadence of 80-90 rpm and task failure was deemed when cadence fell >10 rpm below the targeted cadence, despite strong verbal encouragement. During the final 30 s of baseline bouts and final 30 s of every 5 min throughout cycling trial, 5 single pulse MEPs, 5 SICI paired pulses, 5 LICI paired pulses, and 2 M_max_ were delivered (Figure 1). The order of stimulation type was randomized.

**Figure 1.**
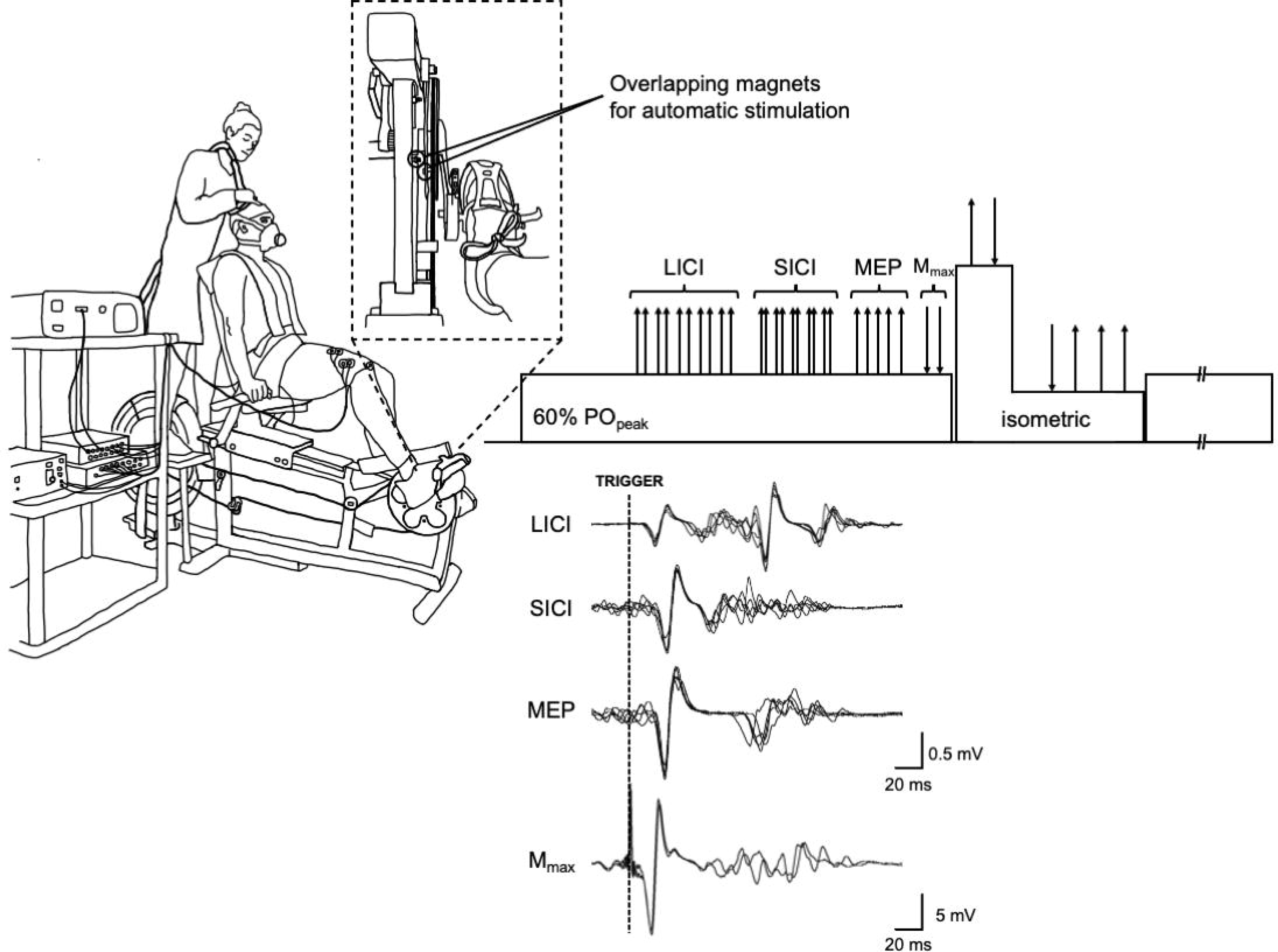
Experimental setup, exercise protocol, and raw signal traces. On a custom-built semi recumbent cycle ergometer with automatic peripheral nerve and transcranial magnetic stimuli, participants performed single-leg cycling at 60% peak power output (PO_peak_) until task failure. Every five minutes, in randomized order, five long interstimulus interval paired pulses (LICI), five short interstimulus interval paired pulses (SICI), five single TMS pulses (MEP), and two femoral nerve stimuli (M_max_) were delivered. Additionally, every five minutes, participants performed a brief maximal and submaximal isometric contraction with stimuli, the results of which are not reported herein.

### Measurements

#### Automatic stimulation setup

A pair of magnetic sensors were positioned on the bike so that one was placed on the medial side of the crank and the other on the trunk of the ergometer setup so that the point at which the magnets crossed would be in the ascending phase of the root mean square (rms) EMG, but nearing the peak, of the upstroke phase of cycling (i.e., 45° crank arm from dead centre). This magnet pairing was setup for automatic M_max_, unconditioned MEP, and SICI stimulations. For LICI stimulations, a third magnet was placed on the crank at a slightly earlier position of the cycling phase so that with the 100 ms interval, the test stimulus would be delivered at the same crank arm angle position as the other stimuli. A trigger signal would be sent once the magnets passed and either of the nerve or TMS setups could be turned on to elicit automatic stimulations. Stimuli were delivered with no delay upon trigger and each stimulation was separated by 5 s.

#### Electromyography

During the experimental visits, EMG was recorded using bipolar Ag/AgCl surface electrode pairs (10 mm diameter, 20 mm interelectrode distance; Kendall MediTrace; Covidien, Mansfield MA) overtop the belly of the vastus lateralis (VL), rectus femoris (RF), and biceps femoris (BF) muscles of the dominant (cycling) leg, with a reference electrode on the patella. At the start of each session, the areas underneath electrodes were shaved and scrubbed with an alcohol wipe prior to placement. Interelectrode impedance of <5 kΩ was confirmed prior to recording, and signals were recorded at a sampling rate of 2000 Hz using PowerLab (16/30-ML880/P, ADInstruments, Dunedin, New Zealand), amplified (ML138, ADInstruments), bandpass filtered (5-500 Hz), and analyzed offline using LabChart 8 software (ADInstruments).

#### Force recordings and peripheral nerve stimulation

Knee extensors force was measured through validated force pedals (LC101-2K, Omegadyne, Sunbury OH) and the signal were amplified (LabChart 8; ADInstruments) and displayed on a monitor in front of participants throughout the entirety of experimental sessions (Doyle-Baker *et al*., 2018). For M_max_, constant-current electrical stimuli were delivered to the femoral nerve of the dominant leg, with the anode (50 X 90 mm; Durastick Plus; DJO Global, Vista CA) on the gluteal fold and the cathode (10 mm diameter; Kendall MediTrace; Covidien, Mansfield MA) atop the femoral nerve in the inguinal fold. While seated and resting in the bike with the dominant leg propped up at the crank angle corresponding to automatic stimulation, single pulses were delivered in increased 10 mA increments (pulse duration: 1000 µs; maximal voltage: 400 V) until reaching M_max_ in both VL and RF muscles. A supramaximal (130%) stimulation was used for the remainder of the session (Visit 1: 96 ± 58 mA; Visit 2: 99 ± 73 mA).

#### Transcranial magnetic stimulation

Prior to determining parameters for TMS, participants performed single-leg cycling for 1 min at the PO corresponding to the session intensity. Without stimulations occurring, the automatic stimulation crank angle markers were used to identify each pedal stroke. Using 10 markers from the start, middle, and end of the 1 min cycling bout, the peak rmsEMG amplitude within a 200 ms window following the marker was obtained and averaged across all 30 pedal strokes. Then, 20% of the averaged maximal cycling EMG from these 30 strokes was set as the isometric contraction intensity for single- and paired-pulse TMS determination.

TMS was delivered to the contralateral motor cortex using 110 mm concave double-cone coils connected to a BiStim unit and two Magstim 200^2^ units (The Magstim Company Ltd, Whitland, UK).

The vertex was first identified as the intersection between nasion to anion and tragi (Aboodarda *et al*., 2019) and marked on a Lycra swimming cap (Aquam Inc., Montreal QC, Canada). For determination of TMS hotspot (Aboodarda *et al*., 2018) and aMT of the unconditioned MEP, participants performed a brief (∼3 s) isometric contraction corresponding to 20% of the averaged maximal cycling EMG at the crank angle corresponding to automatic stimulation. This contraction intensity (∼20% MVC) was chosen so that fatigue induced by multiple contractions required to determine hotspot and aMT could be avoided and so that MEP would not be saturated on the stimulus-response curve (Todd *et al*., 2003). The TMS intensity was then set to 50% maximal stimulator output (MSO) and the coil was adjusted over the left motor cortex (in 1 cm increments) until the greatest motor evoked potential (MEP) amplitude from the VL and RF muscles, and the smallest from the BF muscle, was elicited during a brief contraction. Next, aMT was determined whereby single-pulse TMS was delivered during brief contractions at 20% MVC over the optimal coil site. The aMT determination began at 30% MSO and increased in 2-3% increments, and then 1% increments, until a MEP signal was distinguishable from background EMG. The aMT was confirmed as a consistent MEP response in 3 out of 5 trials, then set to 120% aMT before confirming silent period duration. If silent period was <100 ms (i.e., interstimulus interval duration for LICI), then MSO intensity was increased in 1% increments until the silent period lasted >100 ms (Visit 1: 46 ± 5%; Visit 2: 46 ± 4% MSO).

Paired-pulse stimuli for SICI and LICI was determined during single-leg cycling at the PO corresponding to the session intensity. First, the conditioning paired-pulse stimulus for SICI was determined, as recommended by a recent study (Ruas *et al*., 2020). Briefly, the BiStim was set with a 3 ms interstimulus interval and the predetermined supramaximal aMT test stimulus. Conditioning pulses were delivered at intensities starting from 95% aMT decreasing in 5% increments. The goal was to select a conditioning pulse intensity eliciting a SICI signal that was half the size of the unconditioned MEP; however, this was oftentimes unachievable so the intensity that produced the smallest SICI signal was used (Visit 1: 36 ± 7%; Visit 2: 36 ± 5% MSO). Three paired pulses were delivered at each intensity for verification. This method of SICI determination, rather than using a fixed intensity for all participants, ensures a similar level of inhibition at baseline across the sample (Ruas *et al*., 2020; Zhang *et al*., 2023).

The test paired-pulse stimulus for LICI was then determined. Briefly, the BiStim was set with a 100-ms interstimulus interval and the predetermined supramaximal aMT intensity. Paired pulses with both stimuli set to the same intensity were delivered (Ansdell *et al*., 2020).

### Data and statistical analyses

For VL and RF muscles, the area (from the initial deflection of EMG signal from baseline to the second crossing of the horizontal axis) of unconditioned and conditioned MEPs, as well as the area and peak-to-peak amplitude of M_max_ signals during cycling were identified and measured by visual inspection (Aboodarda *et al*., 2018). Unconditioned MEPs were normalized to the area of M_max_, while SICI and LICI were quantified as the ratio between the average area of five conditioned corresponding pulses over the average of five unconditioned MEP areas. Similar to our prior work (Zhang *et al*., 2021), for each exercise session, variables were linearly interpolated from 0% (i.e., baseline) to 100% (i.e., task failure). The interpolated 20%, 40%, 60%, and 80% values, and raw baseline and task failure values, were used for statistical analysis.

Statistical analyses were computed on absolute values of each variable of interest using IBM SPSS, version 25.0 (IBM Corp., Armonk, NY). Shapiro-Wilk and Mauchly tests assessed the normality and sphericity, respectively, of outcome variables. Where sphericity assumptions were violated, Greenhouse-Geisser was reported. A paired samples t test was used to compare total exercise time between both visits. A two-way repeated measures ANOVA with Bonferroni post hoc analysis was used to compare: i) within-session measures at baseline and ii) between-session measures across six timepoints (raw baseline, 20%, 40%, 60%, 80%, raw task failure). A two-way mixed intra-class correlation coefficient (ICC_3,1_, 95% confidence intervals, and coefficient of variance (CV%) were used to calculate: i) intra-session reliability between both baseline assessments within each condition and ii) inter-session reliability for M_max_ area and amplitude, MEP·M_max_^-1^, LICI, and SICI. ICC_3,1_ was also used to calculate inter-session reliability of all four baseline assessments across both conditions for MEP·M_max_^-1^, LICI, and SICI. ICC values <0.50 were considered ‘poor’, 0.50-0.75 ‘moderate’, 0.75-0.90 ‘good’, and ≥0.90 ‘excellent’ (Koo & Li, 2016).

## RESULTS

Time to task failure was similar in both visits (54 ± 14 min vs. 58 ± 19 min; *p = 0.341*). Upon analysis, SICI and LICI recorded on rectus femoris EMG signals from one male participant were not clearly identifiable and the LICI signals from another two males were unidentifiable (total n = 8 for SICI; n = 6 for LICI reliability analysis in RF muscle).

### Corticospinal measures at baseline

When considering within-session reliability at baseline for corticospinal measures (Table 1, Figure 2), MEP·M_max_^-1^, SICI, and LICI reliability was ranged from good to excellent in both muscles. When pooling and comparing all four baseline bouts (2 baseline × 2 sessions), reliability ranged from moderate to good. When comparing measures at baseline between sessions (Table 1, Figure 3), in the VL muscle, a significant main session effect (F_1,8_ = 6.794, *p = 0.031*) was demonstrated whereby SICI was lower in the second session (0.721 ± 0.113) compared to the first (0.834 ± 0.116). No differences between session were observed for MEP·M_max_^-1^, LICI, or M_max_. In the RF muscle, no differences were noted for any variables.

**Figure 2.**
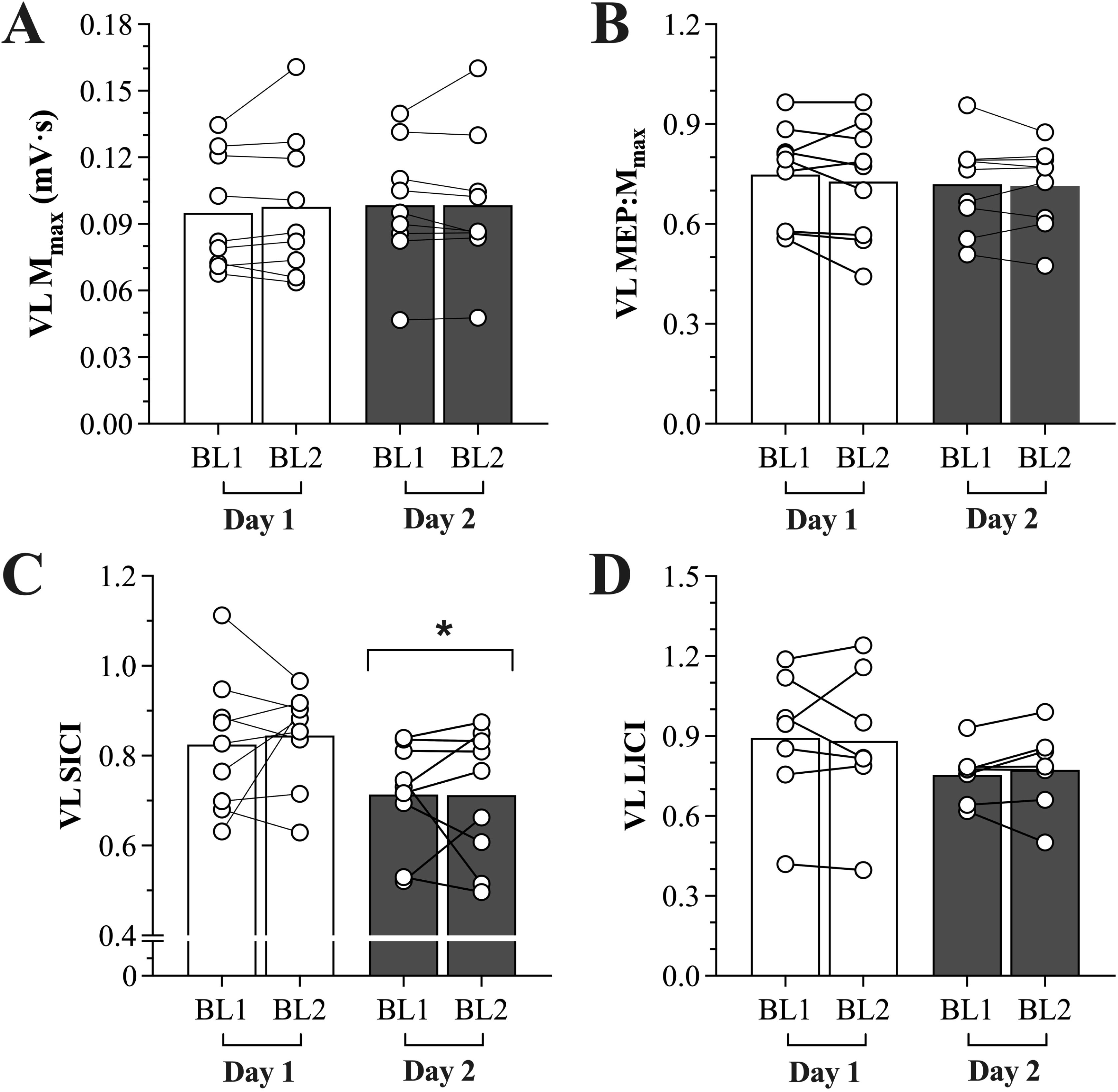
Corticomotor responses of the vastus lateralis at baseline. Maximal M-wave (M_max_) area (A), motor evoked potential (MEP) normalized to Mmax (B), short interval intracortical inhibition (SICI; C), and long interval intracortical inhibition (LICI; D) was measured in two baseline cycling bouts in two separate visits. *difference between visits

**Figure 3.**
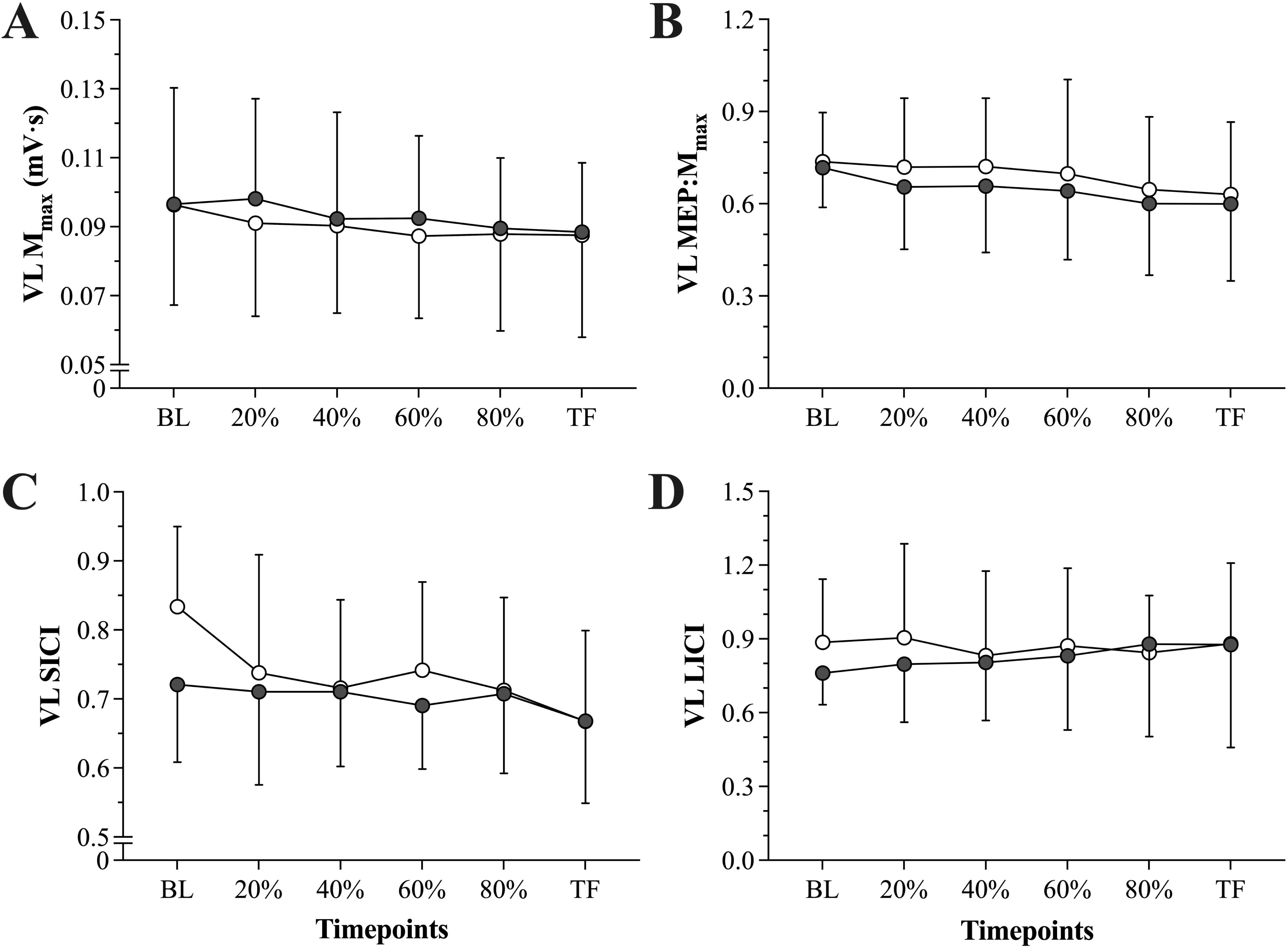
Corticomotor responses of the vastus lateralis across the fatiguing task. Maximal M-wave (M_max_) area (A), motor evoked potential (MEP) normalized to M_max_ (B), short interval intracortical inhibition (SICI; C), and long interval intracortical inhibition (LICI; D) was measured across a cycling task to failure in two separate visits.

**Table 1.** Intraclass correlations (ICC) and coefficient of variation (CV) between corticospinal measures at baseline.

|  |  | ICC (95% CI) | CV (%) |
| --- | --- | --- | --- |
| **VL MEP·M<sub>max</sub><sup>-1</sup>** | Visit 1 | 0.961 (0.841, 0.991) | 21.4 |
|  | Visit 2 | 0.976 (0.894, 0.994) | 17.9 |
|  | All BSLN | 0.894 (0.703, 0.973) | 19.5 |
| **VL SICI** | Visit 1 | 0.750 (-0.161, 0.944) | 15.2 |
|  | Visit 2 | 0.780 (-0.072, 0.951) | 18.1 |
|  | All BSLN | 0.636 (0.125, 0.901) | 18.2 |
| **VL LICI** | Visit 1 | 0.945 (0.672, 0.991) | 28.9 |
|  | Visit 2 | 0.929 (0.621, 0.988) | 16.7 |
|  | All BSLN | 0.803 (0.412, 0.961) | 25.2 |
| **RF MEP·M<sub>max</sub><sup>-1</sup>** | Visit 1 | 0.968 (0.860, 0.993) | 21.5 |
|  | Visit 2 | 0.969 (0.872, 0.993) | 24.7 |
|  | All BSLN | 0.793 (0.458, 0.946) | 24.3 |
| **RF SICI** | Visit 1 | 0.943 (0.716, 0.989) | 24.5 |
|  | Visit 2 | 0.931 (0.642, 0.986) | 33.6 |
|  | All BSLN | 0.827 (0.489, 0.961) | 28.9 |
| **RF LICI** | Visit 1 | 0.818 (-0.557, 0.975) | 31.7 |
|  | Visit 2 | -0.141 (-6.570, 0.839) | 12.6 |
|  | All BSLN | 0.507 (-0.509, 0.919) | 27.6 |

Note: Two baseline measures were performed on each visit and compared, and all four baseline measures across two visits were also compared. BSLN: baseline; CI: confidence interval; LICI: long interval intracortical inhibition; MEP·M<sub>max</sub><sup>-1</sup>: motor evoked potential normalized to the maximal muscle compound M-wave; RF: rectus femoris; SICI: short interval intracortical inhibition; VL: vastus lateralis.

### Corticospinal measures throughout cycling

When comparing between-session measures across the entire exercise task, no significant main condition nor interaction effects were observed for any variables in either VL or RF. In the VL muscle, for MEP·M_max_^-1^, a main effect of time (F_1.5,12.3_ = 4.238, *p = 0.048*) was demonstrated but no pairwise comparisons were significant (Figure 3). For SICI, a main time effect (F_5,40_ = 2.519, *p = 0.024*) was demonstrated but no pairwise comparisons were significant. No significant main nor interaction effects appeared for LICI, M_max_ area, or M_max_ amplitude.

In the RF muscle, for MEP·M_max_^-1^, a main effect of time (F_2.4,18.9_ = 3.526, *p = 0.043*) was demonstrated but no pairwise comparisons were significant. No significant main nor interaction effects appeared for SICI, LICI, M_max_ area, or M_max_ amplitude.

Between-session reliability for MEP·M_max_^-1^, SICI, LICI, M_max_ area, and M_max_ amplitude for both muscles are displayed in Table 2. The reliability of M_max_ area and amplitude was good to excellent in both muscles at all timepoints. In the VL, MEP·M_max_^-1^ ranged from good to excellent reliability, though were considered poor to moderate in the RF muscle. SICI demonstrated poor to moderate reliability for both VL and RF muscles. The reliability of LICI was moderate in the VL at the start of the exercise, but improved to excellent over the course of fatiguing exercise. However, LICI in the RF muscle demonstrated poor reliability.

**Table 2.**
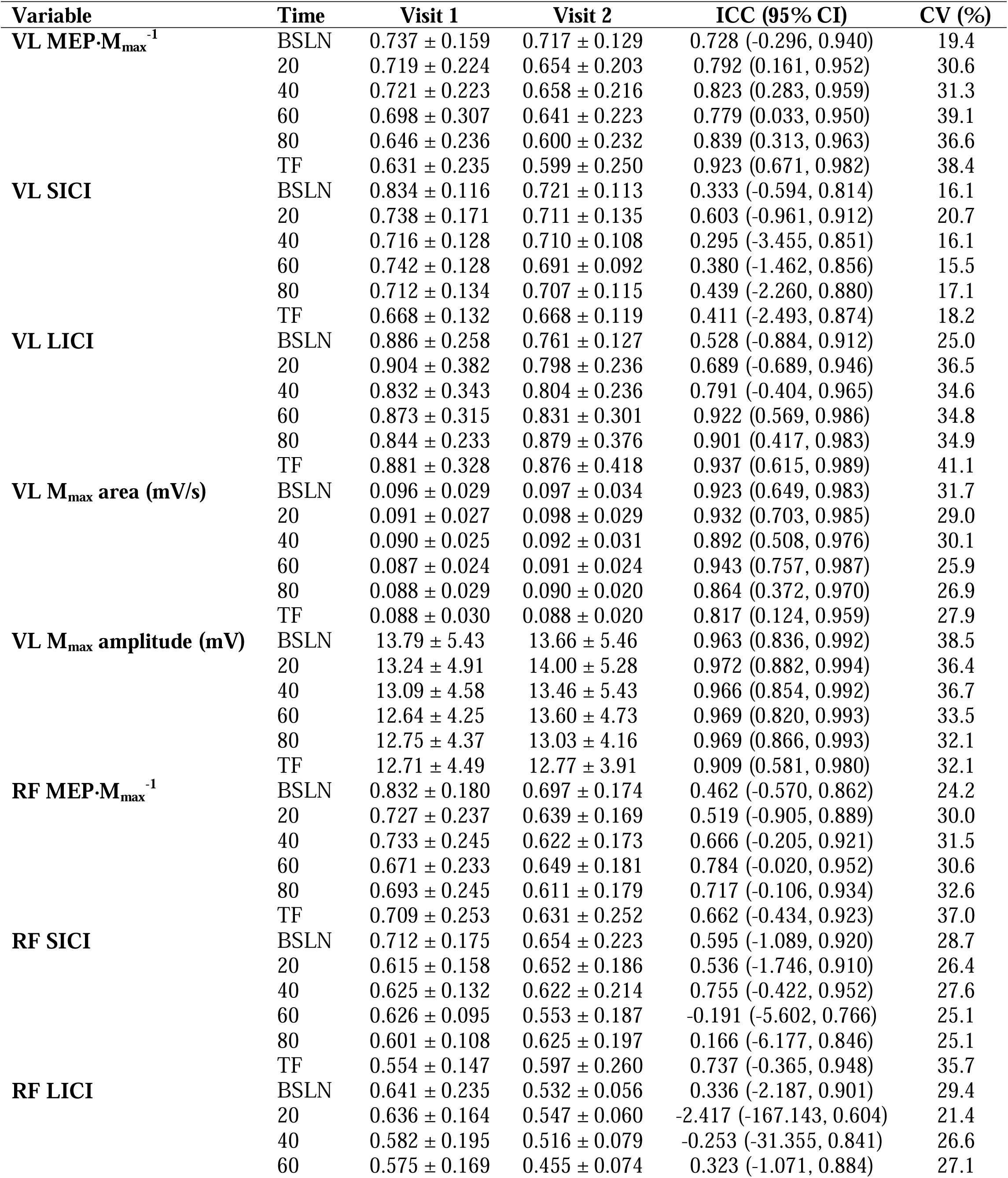
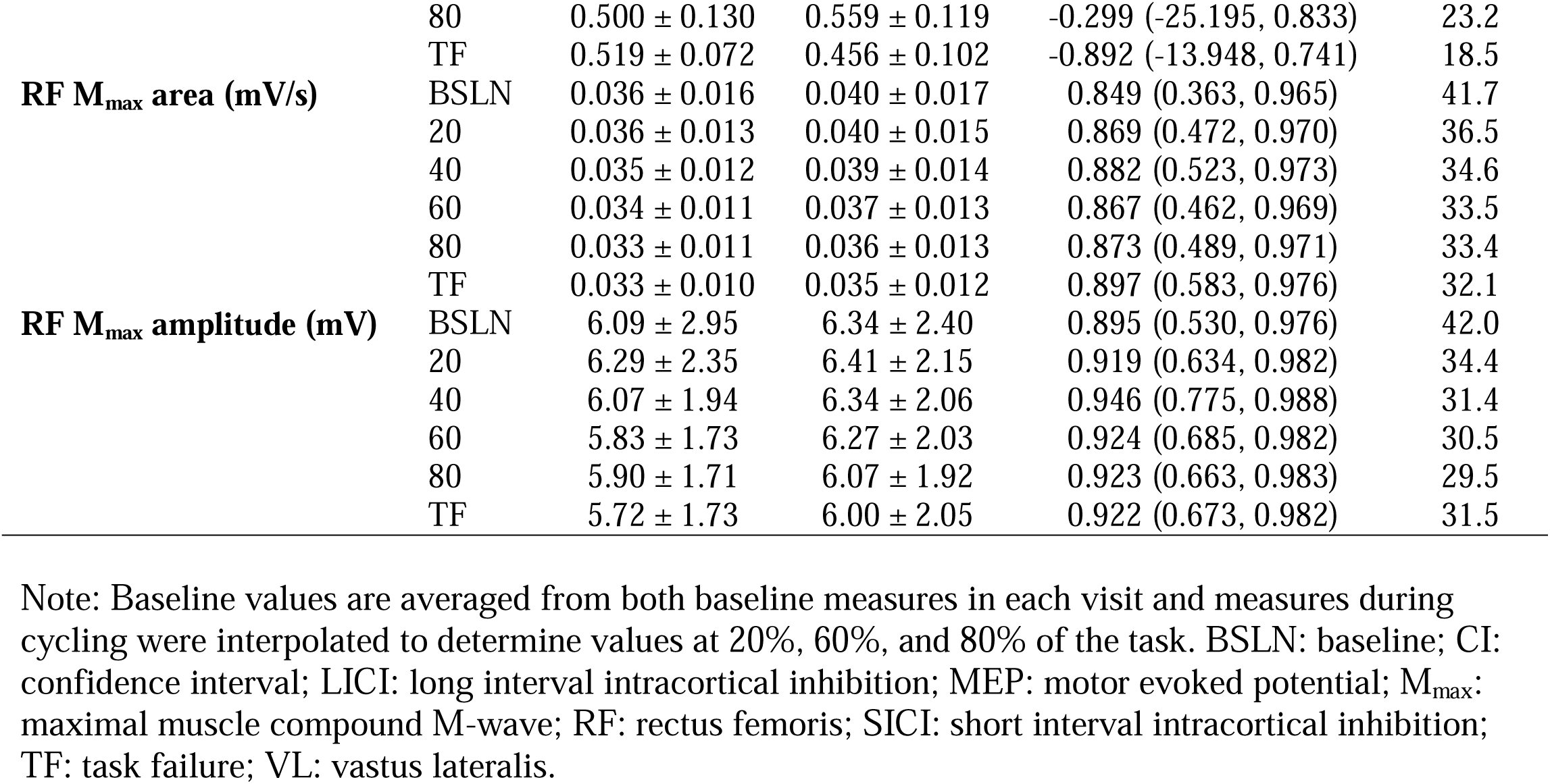
Raw values, intraclass correlations (ICC), and coefficient of variation (CV) of corticospinal measures during cycling.

## DISCUSSION

The present study aimed to assess the reliability of M_max_, MEP·M_max_^-1^, LICI, and SICI measures of the VL and RF muscles during cycling. When comparing the reliability of both baseline bouts *within* the same session, MEP·M_max_^-1^ and LICI were rated excellent while SICI was rated good. At baseline *between* sessions, M_max_ and MEP·M_max_^-1^ demonstrated good reliability, LICI was moderate, and SICI was poor. For the signals recorded throughout the two cycling trials, M_max_ demonstrated consistent and excellent test-retest reliability in both muscles, MEP·M_max_^-1^ ranged from good to excellent, LICI was moderate to excellent, and SICI remained poorly reliable throughout the entirety of the trials. Overall, these results corroborate the cruciality of retaining mode-specific testing measurements and suggest that during cycling, M_max_, MEP·M_max_^-1^, and LICI measures are highly reliable whereas SICI, although demonstrating poor day-to-day reliability, was reliable within the same session.

### Muscle and corticospinal excitability

M_max_ demonstrated excellent reliability within and between sessions, in both non-fatigued (baseline) and fatigued states. This is in line with prior studies that have also observed excellent within- and between- day reliability for M_max_ in quadriceps muscles (O’Leary *et al*., 2015; Latella *et al*., 2017; Leung *et al*., 2018; Brownstein *et al*., 2018). M_max_ has been frequently reported to remain unchanged following various types of fatiguing exercise (Allen *et al*., 2008; Sidhu *et al*., 2018). As such, given that M_max_ regulation (i.e., ionic conductance across the sarcolemma) appears to be relatively stable following most types of exercise excluding long-duration tasks (Krüger *et al*., 2019; Iannetta *et al*., 2022), it can be expected that this measure is highly reliable during cycling, and most variation might be attributed to experimental factors (e.g., electrode movement, automatic stimulation) rather than physiological. Furthermore, the high M_max_ reliability suggests that variation in MEP, LICI, and SICI responses is likely due to supraspinal modulations rather than neuromuscular.

The size of the MEP signal, when normalized to M_max_, represents the functionality and balance of excitatory and inhibitory cortical and motoneuronal influences in voluntary motor command (Weavil & Amann, 2018). Only a few prior studies have measured MEP·M_max_^-1^ during leg cycling (Sidhu *et al*., 2012*b*, 2012*a*, 2017, 2018; Weavil *et al*., 2016), though the reliability of these measures were not reported. Albeit in isometric contractions, MEP·M_max_^-1^ has generally demonstrated strong reliability in lower limb muscles (predominantly VL and RF) across TMS intensities (Luc *et al*., 2014; O’Leary *et al*., 2015; Temesi *et al*., 2017; Brownstein *et al*., 2018; Proessl *et al*., 2021; Pagan *et al*., 2023). For example, Proessl et al. (2021) reported an excellent reliability (ICC = 0.92) and strong variability (CV = 44.3%) of VL MEPs during isometric knee extensions at 15% MVC, an intensity somewhat similar to the present study. Pagan et al. (2023) also recently reported ICC values >0.8 and >0.9 for the MEPs in the VL and RF, respectively, during submaximal contractions in both males and females. In the present study, MEP·M_max_^-1^ demonstrated consistent good to excellent reliability both at baseline and throughout the fatiguing cycling protocol, with variability approximating 20% at baseline and increasing to 30-40% throughout the protocol in both muscles (Tables 1 and 2). Despite the strong reliability, the high variability could be due to increased recruitment of antagonist and accessory muscles, alterations in central motor command, and greater spontaneous fluctuations in cortical and motoneuronal excitation with the development of fatigue (Schilberg *et al*., 2021); this variation should be considered when measuring MEP during cycling and a larger sample size could help reduce variability.

During sustained isometric contractions, MEP·M_max_^-1^ can increase, which may indicate a compensatory mechanism to offset muscle contractile fatigue development (Taylor *et al*., 2016). During cycling, MEP·M_max_^-1^ was not reported to change (Sidhu *et al*., 2012*a*, 2017, 2018; Weavil *et al*., 2016); while the reasons for this discrepancy remain unclear, it emphasizes the importance of utilizing task-specific measurements. It should also be noted that although prior studies have measured spinal motoneuronal excitability (i.e., cervicomedullary or thoracic motor evoked potentials) during cycling, we were not equipped to evoke these signals during the task, so the source of variability in MEP·M_max_^-1^ measures cannot be isolated to the brain or spinal cord. However, our study demonstrates that MEP·M_max_^-1^ during cycling is reproducible and remains so as fatigue develops.

### Intracortical inhibitory circuits

The silent period, a brief window of EMG silence immediately following an unconditioned MEP, is typically measured and represents activity of intracortical inhibitory interneurons (Taylor *et al*., 2006). However, during cycling, identification of a silent period can be difficult to consistently attain due to the configuration of EMG signals during downstrokes. Inhibitory mechanisms can instead be investigated with cortical paired pulse paradigms, as in the present study, or with silent period during an isometric contraction, as traditionally done.

To our knowledge, only one prior study has utilized paired pulse paradigms during cycling phase (Sidhu *et al*., 2018). SICI and LICI, which represent indirect measures of GABA_A_- and GABA_B_-mediated inhibitory circuits (Hanajima *et al*., 1998; Werhahn *et al*., 1999), respectively, can be useful to explore more specific interneuronal cortical mechanisms throughout an exercise task. In the present study, when observing VL muscle responses, SICI demonstrated poor to moderate test-retest reliability (0.3-0.6) across the entire protocol, whereas LICI reliability was moderate for the first part of the task (0.5-0.7) and improved to excellent in the second half (>0.9). This latter observation is intriguing as it can be speculated that homogenized levels of GABA neurotransmitters or GABA_B_ receptor activity within the brain could have reduced LICI variability across the fatiguing task. However, the underlying mechanisms remain unclear and further research is needed to investigate these responses. Additionally, though SICI demonstrated the lowest ICC values out of any of the measures, its CV was the lowest and consistently ranged between 10-20% at baseline and throughout the cycling protocol in the VL, whereas the other variables ranged 20-40% (Tables 1 and 2). This might suggest while the between-individual differences for SICI measurements taken during cycling is large and unreliable, the within-individual precision of measurement may be acceptable.

The modest SICI reliability could potentially be due to the limited number of trials. Prior studies using a fixed conditioning stimulus intensity (i.e., submaximal percentage of aMT) have recommended at least 18 trials for good reliability during isometric knee extension exercise (Brownstein *et al*., 2018).

However, a high number of trials may be impractical to obtain during high-intensity fatiguing exercise. In the study by Ruas et al. (2020) from which we adopted our SICI conditioning stimulus intensity determination approach, the average ICC of different intensities and between sessions was considered poor to moderate. Therein, five paired pulse stimuli were delivered and averaged. Thus, by utilizing a similar intensity approach and number of trials in the present study, we achieved a similar, albeit slightly lower, between-session reliability at baseline (ICC = 0.333 in VL) despite investigating a different contraction mode. Additionally, other studies measuring SICI and LICI of lower limb muscles during isometric contractions have demonstrated equivocal findings. For example, O’Leary et al. (2015) reported moderate to good between-session reliability for SICI and poor reliability for LICI, while Temesi et al. (2017) reported a range of reliability quality (poor to excellent, and even suppressed MEPs) for both SICI and LICI that appeared dependent on the test (conditioned) pulse TMS intensity. Lastly, given that muscle activation increases facilitatory circuit activity and decreases intracortical inhibitory circuits (Ortu *et al*., 2008), this may partly explain the modest reliability of SICI and LICI signals, especially near the beginning of the task (Leung *et al*., 2018).

Of note, for the RF muscle, the assumptions for the ICC model were violated when assessing between- session reliability for SICI at 60%, LICI at 20%, 40%, 80%, and task failure, and when assessing baseline reliability for LICI within Visit 2. This violation was due to negative average covariance between comparisons and produced a negative ICC value, which occurs when the variability within individuals exceeds the variability between (Liljequist *et al*., 2019). This variability could have resulted from the fact that TMS intensity determination for MEP, SICI, and LICI were based on VL responses. Additionally, low statistical power may lead to greater variance in the pattern of LICI across time, which can lead to the negative average covariance observed. While we may expect a relatively similar ICC with a sufficient power given the similarities between both muscles for other reliability comparisons, the sensitivity of this measure should be noted for future considerations.

### Practical applications and methodological considerations

When employing an exercise intervention, it is important to retain the same mode of muscle contraction between the exercise task and the physiological outcomes of interest. This study supports the current evidence that corticomotor responses are mode-specific (Forman *et al*., 2019). Although the paired-pulse measures of cortical inhibition were less reliable, single-pulse MEP and M_max_ were highly reliable even when fatigue incurred. Therefore, to obtain accurate indices of corticospinal excitability during cycling that reflect the true effect of a fatiguing intervention, future studies should take these measures during the same dynamic, rather than isometric, task.

There are some limitations to consider when interpreting the reliability of corticomotor responses during cycling in the present study. As previously mentioned, SP was not measured in this study since the onset and offset of the EMG signal during cycling could not be consistently and accurately detected. Additionally, the mode of exercise was single-leg cycling. As previously mentioned, single-leg cycling was chosen as an framework for a larger investigation that would allow application of experimental pain on the contralateral leg (Aboodarda *et al*., 2019). However, previous studies have postulated that there exists a central nervous system ‘reserve’ during locomotor exercise, which can be accessed during unilateral contractions; in other words, the contralateral hemisphere might contribute to central motor command allowing for interhemispheric facilitation (Lockyer *et al*., 2020). This could factor into the pattern of MEP, SICI, and LICI responses observed across time, or the reliability of such responses.

## CONCLUSION

The present study was the first to measure the reliability of neurophysiological responses during cycling at rest and in fatigued state. This is important for future research studies to employ a methodology that assesses neurophysiological changes in a task-specific manner (e.g., during dynamic exercise). This approach will build on improving historical methods where such measures are obtained during isometric contractions before and after a dynamic exercise, and thus may not accurately reflect corticospinal responses incurred. Additionally, this was the first study to employ paired pulse paradigms to measure cortical inhibition during cycling. Overall, M_max_ and MEP responses are highly reliable in both non-fatigued and fatigued states, while LICI reliability improves with greater fatigue development and SICI signals may not be reproducible across different days with the present single-leg cycling protocol. However, provided participants are not fatigued, the LICI and SICI signals demonstrate good within session reliability. Therefore, future studies can reliably use M_max_ and MEP measures during cycling to evaluate task-specific muscle and corticospinal excitability, respectively, while use of LICI and SICI should be cautioned during fatiguing, heavy intensity single-leg cycling.

